# Herbarium specimens reliably track plant phenological responses to climate change in understudied montane biomes

**DOI:** 10.64898/2026.03.12.709842

**Authors:** Shijia Peng, Brian D. Inouye, Tadeo Ramirez-Parada, Susan J. Mazer, Sydne Record, Aaron M. Ellison, Charles C. Davis

## Abstract

Long-term field observations typically are the “gold-standard” for inferences of phenological sensitivities in montane systems but are spatially limited. Herbarium specimens provide broader spatial coverage, but their utility to accurately capture montane phenology remains poorly known. We compared flowering phenology of 45 species inferred from herbarium specimens with comparable data from nearly 50 years of direct observations at the Rocky Mountain Biological Laboratory. Estimates of flowering time and phenological sensitivity to snow density were consistent between herbarium specimens and observations, but observations revealed secondary flowering peaks.

Herbarium specimens additionally yielded shallower estimates of phenological sensitivity to spring temperature than did field observations. Across co-occurring species, “early” flowering individuals inferred from herbarium specimens, rather than the mean response across all individuals, may better approximate community-level phenological responses to temperature changes. We conclude that herbarium specimens are reliable resources for closing gaps in understanding phenological variation along elevational gradients of montane systems.

## 1. Introduction

Shifts in plant phenology are among the most well-documented biological responses to climatic change (Davis et al. 2015; Park et al. 2018; Thackeray et al. 2016; Willis et al. 2008). Although montane biomes are well-known for their rich species endemism that is highly sensitive to climatic change (Inouye 2020), our understanding of montane plant phenology has been largely restricted to local observational studies (e.g., Inouye 2008; Winkler et al. 2018; Ziello et al. 2009). Because of the complex topography, harsh climate, and scattered populations in montane regions, much less is known about broader-scale geographic and elevational patterns, environmental cues, and underlying mechanisms of montane plant phenology.

Exploring variation in phenological responses among populations and species occupying montane biomes, and especially along elevational gradients, is essential, as such variation may alter ecological and evolutionary processes at a variety of temporal and spatial scales. Differences in phenology among conspecific populations across elevation gradients may alter patterns of gene flow, either facilitating or constraining adaptive evolution through the exchange of locally adapted or maladapted alleles in montane ecosystems (Alleaume-Benharira et al. 2006; Rivest et al. 2003; Sexton et al. 2011; Theobald et al. 2017). In addition to potentially affecting the genetic structure of montane plant species, inter-and intraspecific variation in the timing of flowering in response to warming likely will affect the diversity and seasonal distribution of floral resources (Winkler et al. 2018), the mating system and reproductive success of individual species (Kameyama and Kudo 2009). Climate-induced shifts in flowering time also can alter competitive and facilitative interactions among plant species within montane communities (Klanderud and Totland 2005; Li et al. 2023), especially for species that migrate to higher elevations from lower ones, changing competition for pollinators (Rose-Person et al. 2024; Tiusanen et al. 2020).

Given the challenges of monitoring phenological changes across spatially extensive montane environments, alternative sources of long-term data that assemble and integrate dispersed data sets are crucial for such investigations. A key tool in such efforts are herbarium specimens, which capture snapshots of phenology (e.g., flowering) at specific locations and times, can provide invaluable resources for tracking long-term phenological changes across a vast number of species with extensive geographic and taxonomic coverage (Ahlstrand et al. 2025; Amador et al. 2025; Davis et al. 2015; Park et al. 2024; Ramirez-Parada et al. 2022; Willis et al. 2017). Recent studies have begun to use herbarium specimens to explore montane plant phenology. For example, Munson and Sher (2015) identified that plant species in low elevation habitats in the Southern Rocky Mountains tend to flower earlier in the year than those in high elevation habitats. However, the number of species incorporated, temporal breath, and elevational ranges sampled in these emerging efforts remain relatively limited. Importantly, whether herbarium specimens can faithfully capture montane phenology remains largely untested.

Montane ecosystems are characterized by unique features that could limit the accuracy and generalizability of phenological inferences drawn solely from herbarium data. First, most herbarium sampling is concentrated at relatively accessible lower elevations, whereas remote high-elevation regions are underrepresented (Hughes et al. 2021). Second, botanists often collect specimens at similar times each year (i.e., late spring to summer) out of habit, convenience, or because of other logistical constraints (Schmidt et al. 2025). In montane habitats, collections usually occur long after snowmelt, the timing of which strongly influences the phenology of montane plants (Inouye 2008; Prather, Dalton, barr, et al. 2023). Consequently, average leaf-out or flowering dates recorded from montane herbarium specimens might misrepresent the timing of key phenological events such as the onset, peak, or termination of a phenophase. Third, montane environments are exceptionally variable (Park and Davis 2017); differences in microtopography, slope, and soil conditions that are obscured in larger-scale climatic databases (e.g., PRISM) may influence fine-scale variation in snowpack and temperature (Jaroszynska et al. 2023) and can cause spatial and climatic differences in plant phenology over very short distances. Collectively, these sources of bias raise the question: are herbarium-derived phenological data fully comparable to field observations for exploring phenological responses to climatic change in montane areas?

By comparing phenological data derived from open-access, digitized herbarium specimens with long-term (i.e., 50 years) continuous field observations from the Rocky Mountain Biological Laboratory (RMBL; Figure 1), we tested the validity of using herbarium records to investigate plant phenology under changing climates in the Southern Rocky Mountains, USA, across three dimensions: the distribution of montane plant flowering times, community-level mean flowering responses, and species-specific flowering responses to environmental change. Our approach parallels other validation assessments that have been done for temperate deciduous biomes in the eastern United States (Davis et al. 2015), across the US for species that are well-represented both by herbarium specimens and by field observations using the protocols of the USA National Phenology Network (Ramirez-Parada et al. 2022), and for some tropical regions (Park et al. 2023). We addressed two key questions: 1) how do estimates of changes in phenology and of phenological sensitivity to climatic change in high-elevation montane biomes that are derived from herbarium specimens compare with “gold-standard” estimates from field observations? and 2) can herbarium specimens reliably be used in future studies to explore large-scale elevational patterns in the phenological responses of montane plants to climatic change?

**Figure 1.**
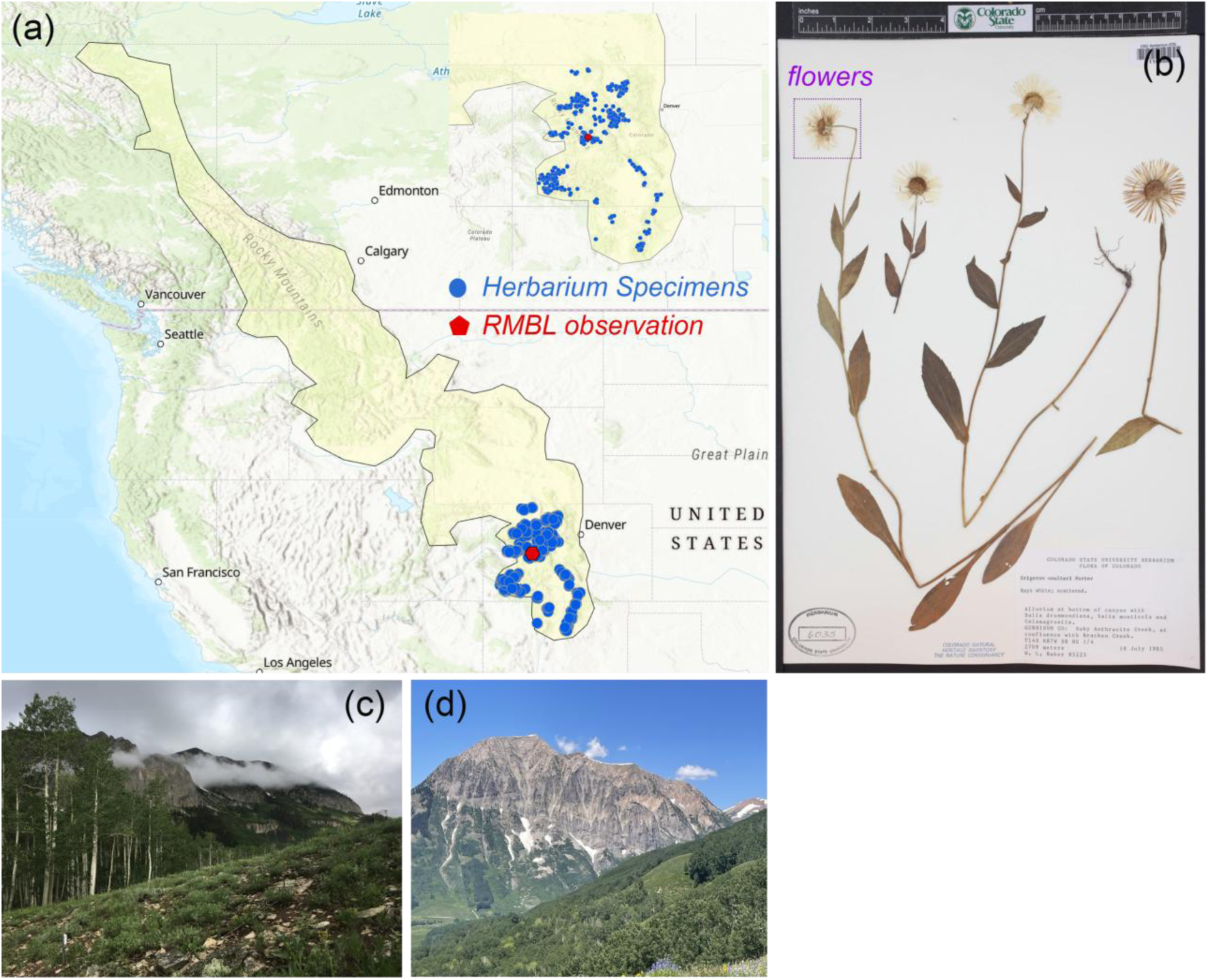
Schematic of the study system. Map of the Rocky Mountain Biological Laboratory (RMBL) location and the collection sites of herbarium specimens included in this study (a). An example of a herbarium specimen of *Erigeron coulteri* (b). Ecoregion of RMBL: sedimentary subalpine forests (c, d).

## 2. Material and Methods

### 2.1 Study location

Our study region, the Southern Rocky Mountains, is among the world’s most well-studied montane areas and has been experiencing very rapid climate change since the mid-1990s (Oldfather et al. 2025; Figure S1 in Appendix S1). The RMBL has been a site for phenology research for over five decades. Relevant studies cover a wide range of taxa and diverse ecological processes, including community structure (e.g., CaraDonna et al. 2014), pollination (e.g., Ogilvie et al. 2017), and phenological shifts under climate change (e.g., Prather, Dalton, barr, et al. 2023). In this subalpine environment, the growing season begins immediately after snowmelt, which is one of the dominant factors influencing flowering phenology (Inouye 2008). Previous studies have suggested that many montane plant species now flower much earlier than in the past because of climate and subsequent earlier snowmelt (Inouye et al. 2008; Prather, Dalton, barr, et al. 2023). Projected climate warming is expected to accelerate snowmelt, reduce snowpack, and potentially intensify drought stress in this region (Lambert et al. 2010; Sheffield and Wood 2008).

### 2.2 RMBL observational and experimental design

Long-term phenological monitoring at RMBL began in 1973 in 35 2 × 2 m plots across meadow and forest habitats. Researchers counted the number of flowers, umbels, or capitula of all species within the plots three times per week during the growing season every year except for 1978 and 1990. We used data from 1975 to 2022 and excluded phenological data from 1973 and 1974 as no environmental data were available for those years. Names, habitats, locations (coordinates for the plot corners), and elevation for each plot are provided in Appendix S1 (Supplemental Methods) and Appendix S2, and are also available in the published dataset (Inouye et al. 2026).

### 2.3 Climate data for RMBL plots

We obtained local climate data from Prather et al. (2023), who integrated daily weather records from five sources in the vicinity of the RMBL: RMBL resident billy barr, the National Oceanic and Atmospheric Administration (NOAA), the United States Geological Survey (USGS), the United States Department of Agriculture (USDA), and Oregon State University’s PRISM Climate Group. The dataset extends from 1975 to 2022 and includes 14 monthly environmental variables including mean temperature, total precipitation, snow density, wind speed, and mean day length (Table S1 in Appendix S1).

### 2.4 Focal species and herbarium specimen collections

Phenological data from herbarium specimens were derived from digitized herbarium specimen images and associated metadata in the SEINet Portal Network, which supports over a dozen regional web portals across North America and includes the Consortium of Southern Rocky Mountain Herbaria (https://www.soroherbaria.org/). We selected species and specimens for analysis for which: 1) there were at least 50 unique collections of the species across space and time in the Southern Rocky Mountains; 2) the specimens included both an exact collection date and location information (i.e., latitude and longitude); and 3) the species had reproductive structures (i.e., buds, flowers and fruits) that were easily identifiable and quantifiable by crowd-workers (see Supplemental Methods). Applying these criteria yielded 29,909 herbarium specimens collected from 1975 to 2022, representing 46 species spanning diverse angiosperm clades for subsequent analysis (Table S2).

We further refined the herbarium observations to include only specimens falling within the same climatic space as RMBL. To do this, we first determined US Environmental Protection Agency (EPA) level IV ecoregion for each herbarium specimen and excluded those were not in sedimentary subalpine forests (the level IV ecoregion of RMBL; https://www.epa.gov/eco-research/ecoregions). Next, we extracted average monthly temperature and precipitation data at a 800-m resolution from PRISM (https://prism.oregonstate.edu/) to characterize the climate of the localities of both the RMBL plots and herbarium specimens. Based on the extracted monthly temperature and precipitation data, we estimated annual mean temperature (abbreviation in WorldClim: bio1), temperature seasonality (bio4), mean temperature of the warmest quarter (bio10), mean temperature of the coldest quarter (bio11), annual precipitation (bio12), precipitation seasonality (bio15), precipitation of the wettest quarter (bio16), and precipitation of the driest quarter (bio17). These variables define a multidimensional climatic space that captures both average and extreme climatic conditions. We pooled these environmental variables for the RMBL observations and herbarium records in a principal component analysis (PCA) and constructed convex hulls in the two-dimensional PCA space separately for RMBL and herbarium observations, retaining only specimens that fell within the RMBL climatic envelope (Figure S2). Last, “*Silene antirrhina”* was excluded because no specimens fell within the same climatic envelope as the RMBL observations. Our final dataset used for direct comparison with RMBL observations included 1,214 herbarium specimens representing 45 species that were collected in a comparable climatic space. We then did crowdsourcing experiments to count the number of flowers contained in each specimen sheet (Supplemental Methods). We excluded herbarium specimens and RMBL observational records in which the number of flowers was zero, leaving 1,120 specimens with records of the number of open flowers on a particular date in the final dataset (median sample size = 24; SD = 13).

### 2.5 Statistical analysis

To assess whether phenological data inferred from herbarium specimens differed from RMBL observations, we first estimated the shape and scale parameters of the Weibull distribution for the flowering day of year (DOY) of each species using maximum likelihood estimation (MLE) with the “*fitdist*” and “*qweibull*” functions in the R package “*fitdistrplus*” (Delignette-Muller and Dutang 2015). Using these parameters, we pooled observations across all years for each species and constructed the cumulative distribution function (CDF) separately for RMBL observations and herbarium specimens. The CDF was defined as 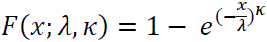, where *x* represents the observed DOY, λ is the scale parameter, and *k* is the shape parameter. We then calculated the DOY corresponding to CDF values of 0.1, 0.5, and 0.9. These quantiles represent the early, middle, and late stages of the flowering period for each species from 1975 to 2022.

To explore whether estimated changes in the flowering DOY determined from the herbarium specimens were correlated with observed changes in flowering DOY observed from RMBL observations under changing climates, we compared the slopes of flowering DOY against key environmental variables such as snow density and spring temperature for both datasets. To obtain monthly snow density for herbarium specimens, we evaluated three regression algorithms (Supplemental Methods) and selected a random forest (RF) model with the best fit (Figure S3) to predict snow density from three climate variables (temperature, precipitation, and temperature range) using data from Prather et al. (2023). These variables were first identified in a RF model as being the most important uncorrelated predictors of snow density (r < 0.7; Table S3). Temperature range (i.e., the difference between monthly maximum and minimum temperatures), reflects seasonal temperature variability that may influence snowpack compaction (Hamlet et al. 2005). We then applied this RF model to monthly temperature and precipitation data from PRISM to predict a monthly snow density value for each herbarium specimen on the date when it was collected.

To evaluate the reliability of PRISM data in representing the climatic conditions of RMBL observations, we additionally matched PRISM climatic data with RMBL observations and applied RF to predict snow density for each observation. The median predicted snow density across models with high predictive accuracy (*R^2^* > 0.6; Supplemental Methods) was used as the final estimates. This approach allowed us to estimate snow conditions specific to the collection date and location of each specimen, providing a more consistent climatic context for comparison with observational records.

Previous studies suggest that the seasonal montane habitat in the Southern Rocky Mountains typically has a persistent snowpack from November through May (Prather, Dalton, barr, et al. 2023). At RMBL, the estimated mean snowmelt date from 1975 to 2022 occurred on DOY 139 (i.e., mid-May) based on data from Prather et al. (2023) (Figure S4). To assess the impact of snow conditions on flowering phenology, we developed three separate models relating flowering DOY to three snow density variables: 1) annual mean snow density; 2) mean snow density in March and April and 3) snow density in May. We constructed these three models for RMBL observational records with local climatic data from Prather et al. (2023).

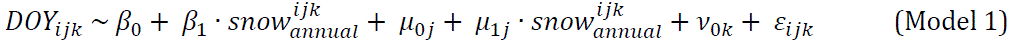

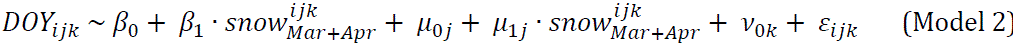

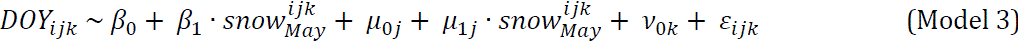

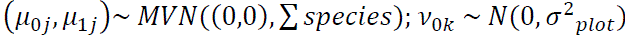

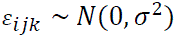

where *DOY*_*ijk*_ represented flowering DOY for the *ith* observation record of species *j* in plot *k*, which was equal to the intercept term β_0_ plus the fixed effects of snow density β_1_ and random effects μ_0*j*_ and μ_1*j*_. The random intercept μ_0*j*_ allowed different species to have different mean flowering DOY, while the random slope μ_1*j*_ accounted for species-specific responses to snow density. The random intercept ν_0*k*_ represented plot-level variation. ε_*ijk*_ was an error term. Each of the three models used the same structure but substituted the snow density with one of three forms corresponding to the year of RMBL observation. All models were constructed using linear mixed models (LMMs) with the “*lmer*” function in the “*lmerTest*” package (Kuznetsova et al. 2017).

We separated snow density variables because different periods of snow accumulation may have distinct effects on flowering DOY. For example, late-winter snowpack can influence early season soil condition and bud development, whereas snowmelt in May may determine the onset of the growing season that could directly affect flowering timing. We then calculated the Akaike Information Criterion (AIC) for each of these three models and selected the snow density variable that yielded the lowest AIC as the best predictor variable.

We also used LMMs to test whether the slope of flowering DOY in response to the selected snow density variable differed among the three datasets. For the full model, flowering DOY was the response variable, snow density, the type of dataset, and their interaction were included as fixed effects. The three datasets were: herbarium specimens with PRISM climatic data, RMBL observational records with local climatic data from Prather et al. (2023), and RMBL observational records with PRISM climatic data. The number of flowers for each species was also considered as a covariate since flower number indirectly indicates plant vigor, health, and size, all of which can influence flowering date (i.e., larger, healthier individuals may delay flowering because they may spend more time acquiring resources during the vegetative stage). For RMBL observations, we calculated the maximum number of flowers recorded for each species in each year because RMBL observations track repeated flowering counts of the same population throughout a season. Therefore, the annual maximum flower number can best represent the potential reproductive output of each species in a given year. In contrast, herbarium specimens represent single, opportunistic collection events of individual plants. For these specimens, we used the number of flowers counted on each individual, which reflects its reproductive state at the time of collection. Species were included as a random intercept, and random slopes of interaction between snow density and dataset type were specified for each species to account for species-specific variation in phenological responses to snow density estimated from different types of datasets. We also included a random intercept for plot; RMBL observations retained their plot identifier but herbarium specimens were assigned a single “NA” to allow inclusion in the model.

The significance of the predictor variables in models was evaluated using the *z* distribution to obtain *P*-values from the Wald *t*-values provided by the model output (Luke 2017). Differences among species in their relative sample sizes in each dataset may result in different contributions to the cross-species mean slope, which can generate differences in the estimated slopes among different data sources. We thus additionally extracted the species-specific slopes within each dataset and examined how closely these paired estimates clustered around the 1:1 line. We used paired *t*-tests to examine whether the mean difference between species-specific slopes from each of the two datasets differed significantly from zero. We also calculated Pearson’s correlation coefficients to assess whether the herbarium specimens provide a reliable predictor of species-level phenological responses to environmental change.

We further used LMMs to examine whether flowering DOY responded consistently to spring temperature across the three datasets. Here, we defined spring temperature as the mean temperature from April to June, which has been identified as the most relevant period influencing flowering phenology and captures post-snowmelt thermal conditions at RMBL (Diez et al. 2012; Inouye 2008). All model structures were the same as those used for snow density.

Since flowering DOY measurements across consecutive years for the same individuals are not independent, we adopted two alternative approaches to examine whether non-independence among repeated observations affected the results (Supplemental Methods).

## 3. Results

### 3.1 Comparison of flowering phenology distributions derived from RMBL observations and herbarium specimens

We reported the shape (*k*) and scale (λ) parameters of the Weibull distribution fit to flowering DOY for each species in Table S4. Overall, early, median, and late flowering times inferred from herbarium specimens were generally consistent with those from RMBL observations across species. The estimated mean early flowering time among species was approximately DOY 170, representing the period during which the earliest 10% of individuals flowered (Figure 2a, d). The median flowering time (50% quantile) occurred on approximately DOY 200 (Figure 2b, e), and the late flowering time (90% quantile) was approximately DOY 220 (Figure 2c, f). However, the range of variation in flowering time among species was larger in the RMBL observations than in the herbarium specimens. The RMBL observations showed secondary community-level peaks in the number of species flowering, with local surges of species in early flowering between DOYs 130-150, surges between DOYs 160-180 for middle flowering, and surges between DOYs 180-200 for late flowering.

**Figure 2.**
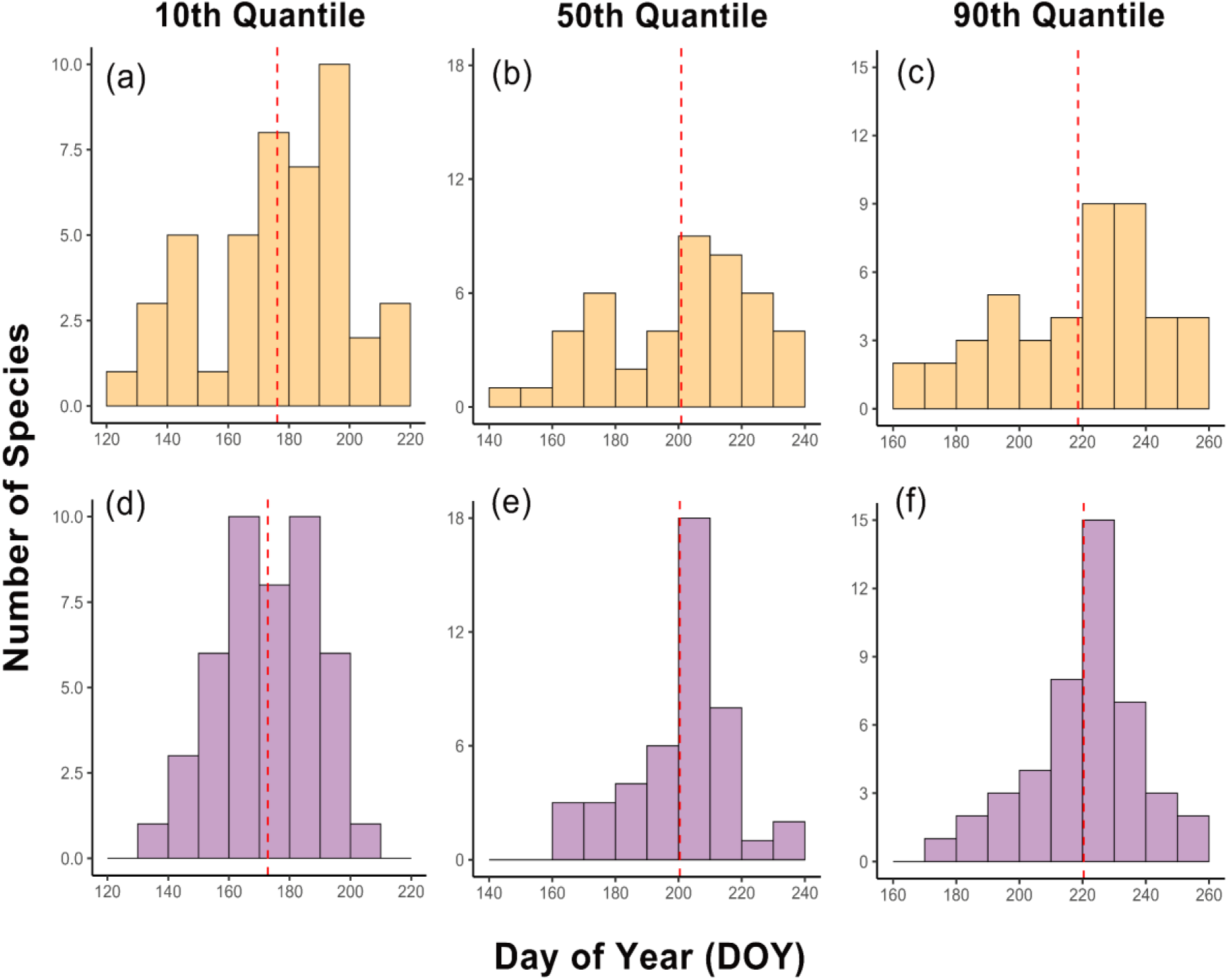
Histograms of the estimated 10th, 50th, and 90th quantiles of flowering day of year (DOY) across all species. The early-season (10th quantile), peak-season (50th quantile), and late-season (90th quantile) flowering DOY for each species were estimated using Weibull models fitted to RMBL field observtaions (a–c) and herbarium specimens (d–f).

### 3.2 Phenological sensitivity to snow density

Models using snow density in May yielded lower AIC values (AIC = 8,627.34) than those with annual mean (Δ*AIC* = 1576.37) or March–April mean (Δ*AIC* = 1532.17) snow density. In general, higher snow density in May and a larger maximum number of flowers were associated with later estimated and observed flowering DOY across all species. However, there was no significant interaction between snow density and dataset type (*p* > 0.05; Table S5). The estimated among-species mean slopes of flowering DOY in response to snow density did not differ among herbarium specimens and RMBL observations (*p* > 0.05 for paired *t*-test; Figure 3a; Figure 4a-c). Species-specific slopes obtained from RMBL observations and those derived from herbarium specimens were moderately positively correlated (r = 0.5; Figure 4b-c). In turn, species-specific slopes derived from RMBL observations but obtained using different climate datasets were highly positively correlated (r = 0.93; Figure 4a).

**Figure 3.**
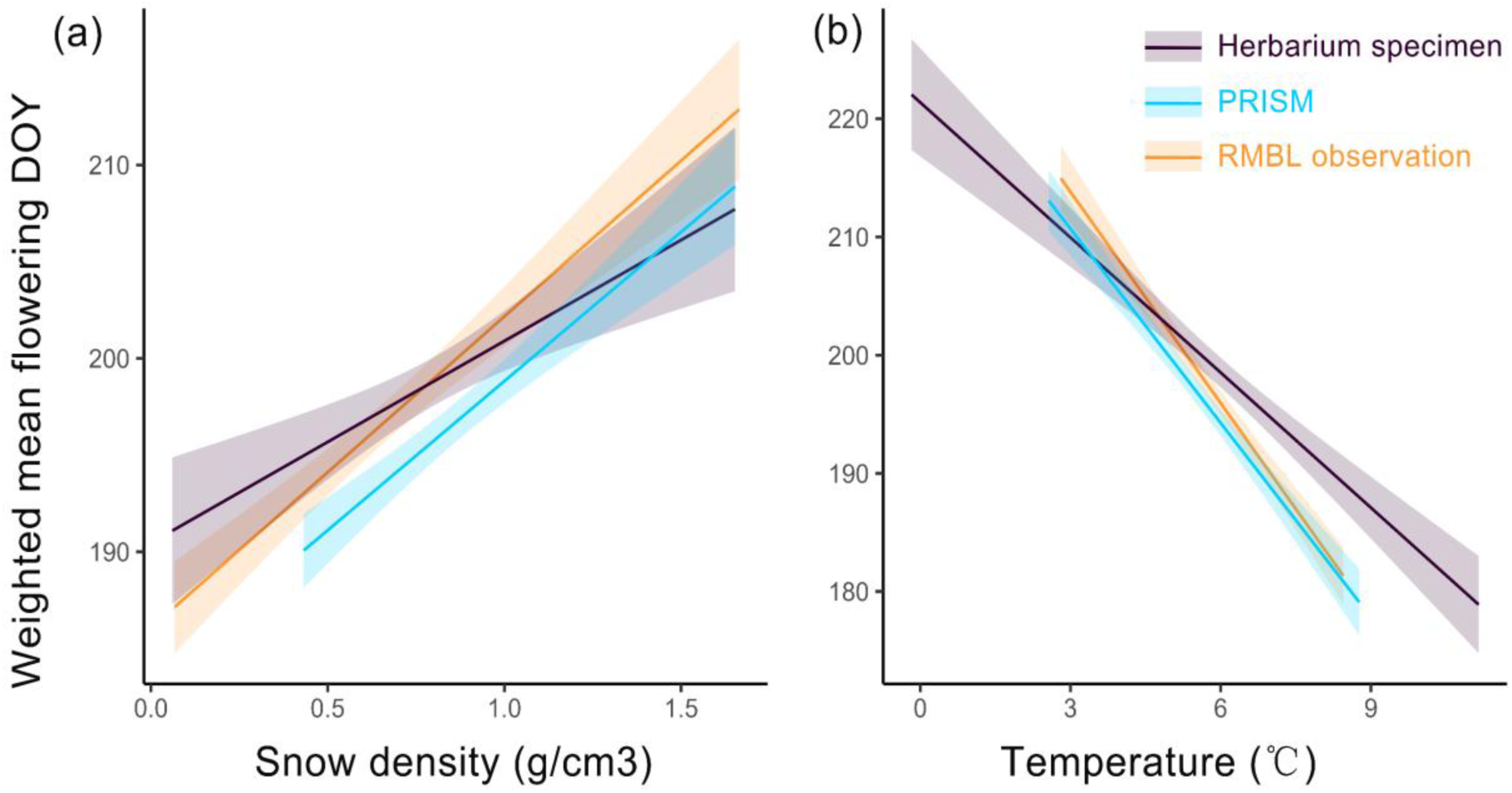
Effects of snow density in May (a) and spring mean temperature (from April to June, b) on flowering day of year (DOY) within the RMBL climatic space estimated based on different databases. Purple lines represent relationships estimated from herbarium specimens with PRISM monthly climate data; light blue lines represent relationships estimated from RMBL field observations with PRISM monthly climate data, and orange lines represent relationships estimated from RMBL field observations with local climate data from Prather et al. (2023). Shaded areas indicate 95% confidence intervals.

**Figure 4.**
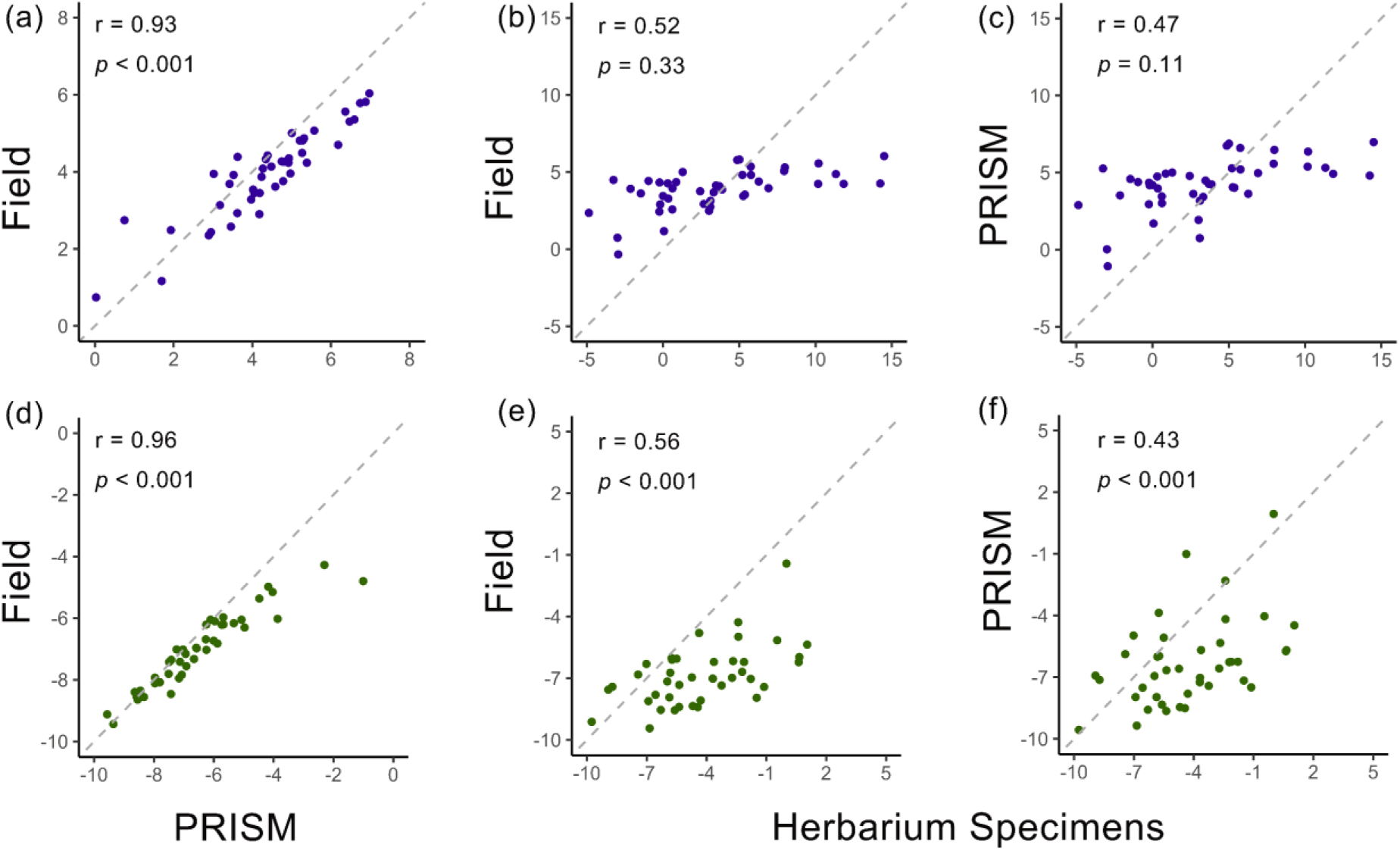
Comparisons of phenological responses to snow density in May (purple points; a–c) and spring mean temperature (April–June; green points; d–f) between herbarium specimens and RMBL field observations. Each point represents the species-specific slope of the relationship between flowering time and snow density (or spring temperature). Flowering time was quantified as day of year (DOY). Each panel shows pairwise comparisons among three data sources: RMBL field observations with local climate data from Prather et al. (2023) versus RMBL field observations with PRISM climate data (a, d); herbarium specimens with PRISM climate data versus RMBL field observations with local climate data from Prather et al. (2023) (b, e); and herbarium specimens with PRISM climate data versus RMBL field observations with PRISM climate data (c, f). Grey dashed lines represent the 1:1 relationship.

Quantile regressions further showed that the among-species mean slopes of the 10th, 50th, and 90th quantile flowering DOY related to snow density, estimated from herbarium specimens, were not significantly different from the 50th quantile slope estimated from RMBL observations (all *p* > 0.05 for paired *t*-test; Figure 5a-c). However, the species-specific slopes showed a relatively low correlation between the two datasets (r ≈ 0.25).

**Figure 5.**
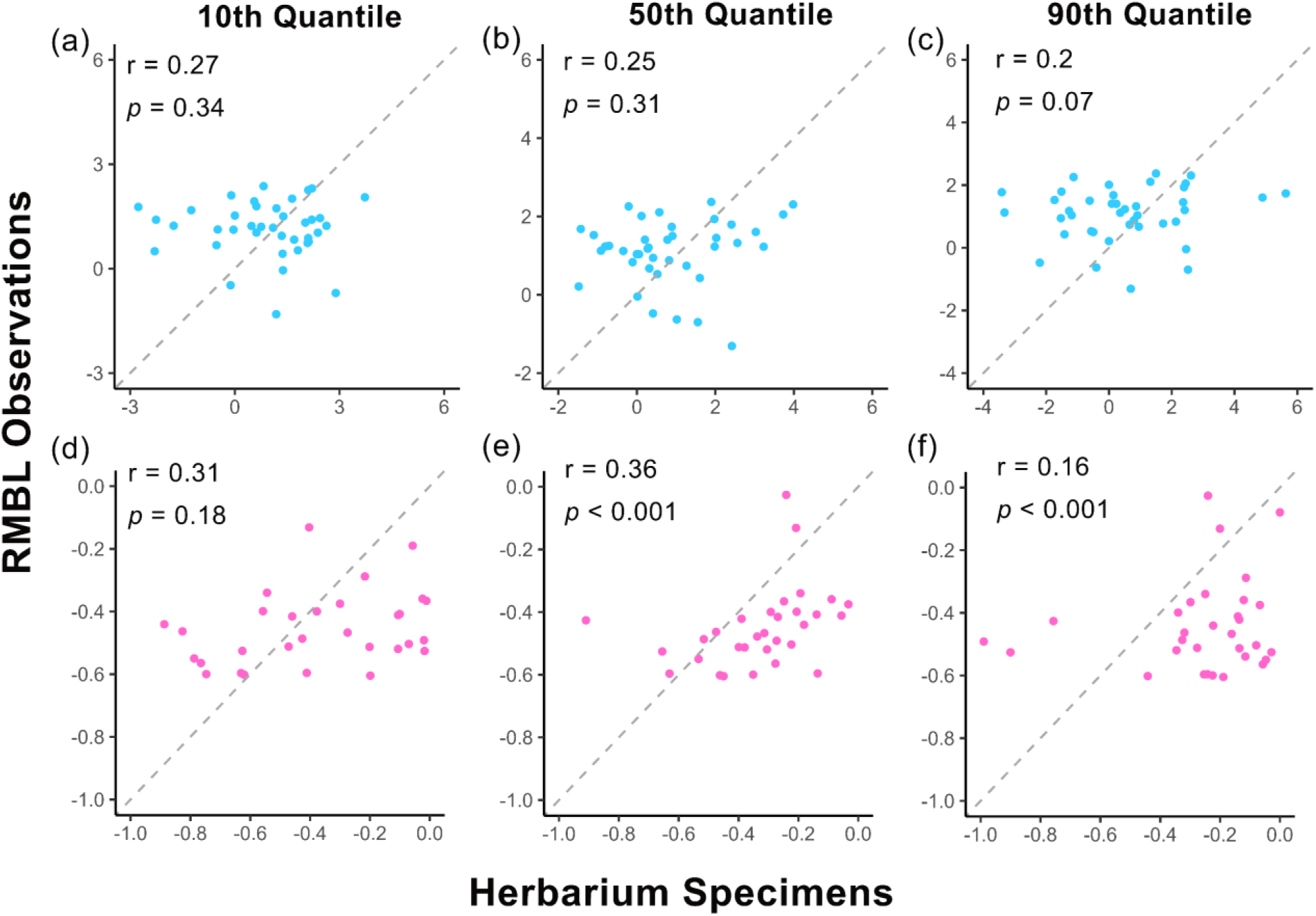
Quantile regression comparisons of phenological responses to snow density in May (light blue points; a–c) and spring mean temperature (April–June; pink points; d–f) between herbarium specimens and RMBL field observations. The 10th (a, d), 50th (b, e), and 90th (c, f) quantile regression slopes of herbarium specimens with PRISM climate data were compared with the 50th quantile regression slopes of RMBL local observations with local climate data from Prather et al. (2023). Flowering time was quantified as day of year (DOY). Each point represents the species-specific quantile slope of the relationship between flowering DOY and snow density (or spring temperature). Grey dashed lines represent the 1:1 relationship.

### 3.3 Phenological sensitivity to spring temperature

Generally, high spring temperature (i.e., mean temperature from April to June) and a small maximum number of flowers advanced flowering DOY. There was a significant interaction between spring temperature and dataset type (*p* < 0.001; Table S6). The average slope of flowering DOY in response to spring temperature was significantly shallower for herbarium specimens than for RMBL observations, whether using PRISM or RMBL-observed local climatic data (*p* < 0.001 for paired *t*-test; Figure 3b; Figure 4d–f). Species-specific slopes exhibited a moderate correlation between herbarium specimens and RMBL observations (r = 0.5; Figure 4e-f). Similarly, the average temperature sensitivity was significantly shallower for RMBL observations with PRISM climatic data compared with those with local climatic data, but the species-specific slopes show a high correlation between two datasets (r = 0.96; Figure 4d).

Quantile regressions further showed that the mean slopes of 10% quantile flowering DOY in response to spring temperature, estimated from herbarium specimens, were most similar to the 50% quantile mean slopes estimated from RMBL observations (*p* = 0.181 for paired *t*-test; Figure 5d). However, the slopes of the 50% and 90% quantile flowering DOY from herbarium specimens were shallower than the 50% quantile slopes estimated from RMBL observations (Figure 5e, f). The species-specific slopes showed a relatively low correlation between the two datasets (r ≈ 0.36).

Two alternative approaches show that using the mean flowering DOY of each species in each year for RMBL observations and incorporating random slopes for year × dataset type interactions did not change parameter estimates in either the temperature or snow density model, suggesting the main results were robust to potential influences of year to year variation and to data non-independence (Table S7–S10).

## 4. Discussion

### 4.1 Herbarium specimens generally provide reliable estimates of flowering time for montane plants

Herbarium specimens provide biased, unevenly sampled (among species, years, space), and incomplete records of phenological events (e.g., Daru et al. 2018; Davis et al. 2015; Schmidt et al. 2025; Willis et al. 2017). From comparisons between herbarium specimens and RMBL observations in their distribution of flowering times, community-level mean responses, and species-specific responses to environmental changes, our results demonstrate that with approximate caveats, herbarium specimens generally can serve as useful proxies for studies of plant phenology in montane areas. However, caution is warranted when interpreting species’ phenological responses to climatic change, as their agreement with field estimates depends on the type of climatic drivers, biological scale, and flowering stage under consideration.

There was good agreement in mean observed flowering times of montane plants inferred from digitized herbarium specimens. However, secondary community-level peaks in flowering time across all flowering stages were observed only in the RMBL observations. Thus, herbarium data may mask more dynamic phenological variability within a season when compared to observational data. For example, the among-species mean early-flowering time (10^th^ quantile) estimated from herbarium specimens occurred on DOY 173, whereas field observations revealed an earlier (and smaller) peak of early-flowering among species between DOYs 130-150 in addition to a primary peak at DOY 176. This earlier peak in flowering may reflect the rapid onset of the growing season following snowmelt at RMBL (Inouye 2008) when collections are reduced or absent due to harsh field conditions or the inconspicuousness of early-flowering individuals. Similarly, during mid-flowering (i.e., 50^th^ quantile), field observations demonstrated an earlier secondary flowering peak between DOY 160-180, corresponding to the among-species mean flowering time in the “early-flowering” herbarium specimens. This result supports the conclusion that herbarium specimens are generally collected in a narrower time window (*sensu* Davis et al. 2015) and may incorrectly compress flowering times for early-flowering species.

### 4.2 Effects of snow density on flowering time of montane plants

We did not find significant differences in the estimated average responses of flowering DOY to snow density in May between herbarium specimens and RMBL observations. The timing of snowmelt is a primary determinant of flowering time for most montane plant species, with flowers blooming immediately after snowmelt (Forrest et al. 2010; Inouye 2008; Miller-Rushing and Inouye 2009). Climatic warming is leading to earlier snowmelt, which, in turn, normally advances the onset of the growing season and shifts flowering time forward (Jabis et al. 2020; Rose-Person et al. 2024; Winkler et al. 2008). Our quantile regression results demonstrate that, on average, the slopes of the 10th, 50th, and 90th quantiles of flowering DOY against snow density, estimated from herbarium specimens, are consistent with the slope of the 50th quantile DOY derived from RMBL observations. Therefore, even though flowering times recorded in herbarium specimens are generally later than those directly observed in the field, both herbarium specimens and field observations capture a consistent, community-level response to snowmelt across the season, and provide comparable estimates of average phenological sensitivity to snow density among species within a community.

### 4.3 Effects of spring temperature on flowering time of montane plants

The slope of the average responses of flowering DOY to spring temperature from herbarium specimens is significantly shallower than that obtained from RMBL observations. This suggests that herbarium specimens provide more conservative (i.e., lower) estimates of phenological sensitivity to spring temperature than do field observations. Montane plants often are highly sensitive to early post-snowmelt temperatures, as heat accumulation during this period directly triggers the onset of growth and development of flower buds (Jabis et al. 2020; Livensperger et al. 2016). Our analysis of flowering time distributions of montane plants supports the idea that herbarium specimen collections often are biased towards individuals at or near peak (or late) flowering, when floral displays are most conspicuous (cf. Davis et al. 2015; Panchen et al. 2019; Schmidt-Lebuhn et al. 2013). Herbarium specimens thus tend to miss early flowering individuals, primarily capturing the average flowering time of a population rather than the onset of flowering. As a result, the effects of early post-snowmelt temperature variation on flowering time may be smoothed out in herbarium specimens, leading to more conservative estimates of the slope in the relationship between flowering time and spring temperature. Conversely, field observations can capture substantial shifts in flowering time even under conditions of relatively small temperature fluctuations during the early post-snowmelt period. Previous phenology studies based on herbarium specimens generally characterized flowering DOY using the mean value (e.g., Park et al. 2018; 2022; Primack et al. 2004), our results suggest that in montane systems, herbarium specimens of relatively “early” flowering individuals may provide the best approximation of the population’s average flowering time.

Notably, both the geographic extent and elevational ranges encompassed by the herbarium specimens we included were still broader than those covered by RMBL observations. Field observations typically monitor populations growing under relatively homogeneous environmental conditions at RMBL sites (∼ 2900m), whereas herbarium specimens include individuals collected across a broader range of elevations (Figure S5; ≈2434–3755m). Studies suggest that populations at different elevations inhabit distinct microenvironmental conditions and may exhibit different local adaptations to snowmelt, even within the same macroclimatic zones (e.g., Winkler et al. 2018). Such variation can influence the phenotype of adaptive traits, including tolerance to chilling and photoinhibition, ability to develop under snow, and seed dormancy (Bemmels and Anderson 2019; Germino and Smith 2000; Keller and Körner 2003). An alpine transplant experiment with *Ranunculus acris* in Western Norway, for example, demonstrated that both phenotypic plasticity and genetic adaptation affect the phenological response to snowmelt, and individuals from high-elevation and wet sites showed limited response to advanced snowmelt (Delnevo et al. 2018). Therefore, datasets spanning broad elevational gradients may produce slope estimates that integrate responses across heterogeneous local environments, contributing to the relatively lower cross-dataset correlation in species-specific slopes.

### 4.4 Limitations and paths forward

Our study has two limitations. First, we only compared the mean phenological responses between herbarium specimens and RMBL observations. Since specimen availability is limited in a given year, we were unable to robustly estimate phenological parameters beyond mean flowering DOY such as onset, termination and duration of flowering, which restricts comparisons with field observations of other phenological stages. Unlike observational studies that include repeated observations of the same individuals, herbarium specimens typically record the phenological status of a randomly selected individual on a specific date and year. A collected individual could be among the earliest-or latest-flowering in the population, or anywhere in between. Quantile regression may serve as a complementary approach for characterizing phenological shifts across flowering stages and for reducing uncertainty on estimates of mean flowering response. Although herbarium specimens span greater temporal and spatial scales than field observations and may complement the results from field observations, they are not a replacement for regularly censused, continuous observational investigations (Park et al. 2023).

Second, our study included only 45 montane plant species, because inferring phenophases from herbarium specimens can be challenging for some taxa. For instance, some representative plant species of *Salix*, *Carex*, or *Polygonum* have tiny flowers that are often aggregated or overlapping within specimens and are difficult for crowd-workers to count accurately. With the rapid development of computer vision leveraging artificial intelligence and machine learning, it is becoming increasingly feasible to automatically extract phenological data from digitized specimens. These approaches may make it possible to extract phenological information even for taxonomic groups with small, clustered, or overlapping reproductive structures (e.g., Love et al. 2021; Triki et al. 2021), although their wide application still requires careful optimization and quality control.

## 5. Conclusions

In summary, due to the unique environments of montane biomes, herbarium specimens underrepresent early flowering individuals and species, estimate lower phenological sensitivity to spring temperatures, and, in our samples, do not document secondary (earlier) community-level flowering peaks. “Early” flowering individuals of herbarium specimens may best approximate a population’s average flowering response to spring temperature. Despite these caveats, herbarium specimens remain a valuable, and indeed often the only, resource for addressing phenological knowledge gaps in understudied montane regions. The next challenge lies in scaling up our assessments to capture phenological responses across broader elevational gradients and diverse climatic contexts to deepen our understanding of how montane plant phenology responds to future climate change.

## Acknowledgements

We acknowledge funding from Harvard University and from National Science Foundation funding grants (CC Davis): DEB-1754584, EF-1208835, DEB-2101884, DEB-1802209, and MRA-2105903. Additionally, for BD Inouye, the projects supporting phenology data collection include DEB-0238331, DEB-0922080, DEB-1354104, DEB-1912006, and DEB-2016749.

## Author Contributions

SP, AME and CCD conceived the idea. SP designed the project. BDI provided RMBL field data. SP led crowdsourcing experiments. SP analyzed and visualized the data with guidance from AME, BDI, CCD, TRP, SJM and SR. SP drafted the initial manuscript. AME, BDI, TRP, SJM, SR and CCD reviewed and edited the manuscript.

## Data Availability Statement

The original phenological data and analysis code used to reproduce the results of this manuscript are publicly available. The data are deposited in a Zenodo repository (doi: https://doi.org/10.5281/zenodo.18866958) and are also available on GitHub (https://github.com/Shijia818/Rocky_Mountain_validation).

## Supplemental Methods

### RMBL observational and experimental design

Long-term phenological monitoring at RMBL began in 1973, when D. Inouye and his colleagues established a series of 2 × 2 m plots across meadow and forest habitats including the original Rocky Meadow plots (plot codes: RM1–RM7) and aspen forest (RM8–RM9) to track flowering dynamics. Additional plots were added in 1974 in wet meadows (WM1–WM5), willow-meadow interfaces (INT1–INT5), and *Erythronium* meadows (EM1–EM2) to capture broader ecological variation. Experimental treatments, including *Veratrum* removal plots (VR1–VR2) and open-topped “greenhouse” warming plots (GH1–GH5), were later added to examine species interactions and global warming effects. More recently, four plots were established on a sloped meadow (SG1–SG4), and two plots were added in new habitats (MDW established in 2004–2012; MDW2 in 2013–2022, and stream plot STR in 2004). Researchers counted the number of flowers, umbels, or capitula of all species within the plots three times per week during the growing season every year except for 1978 and 1990. We used data from 1975 to 2022 and excluded phenological data from 1973 and 1974 as no environmental data were available for those years. Coordinates for the plot corners, and elevation for each plot are provided in Appendix S2, and additional detail at https://osf.io/p9vuj.

### Crowdsourcing experiments and phenological data extraction

Crowd-workers hired through Amazon’s Mechanical Turk service (MTurk; https://www.mturk.com/) counted the number of flowers contained in each specimen sheet using the CrowdCurio interface (Willis et al. 2017). Crowd-workers were required to achieve at least 80% accuracy across two test trials in identifying and quantifying flowers (or inflorescences) before they could proceed to actual data collection (Park et al. 2018). To estimate the reliability of each crowd-worker, each image set scored by a single crowd-worker included nine unique images and one duplicate image randomly selected from the other nine (Park et al. 2018; 2022). We calculated the reliability score for each crowd-worker as 1 minus the absolute difference in flower counts between two duplicate specimens, divided by the total flower count across both specimens (1– |count1-count2|/|count1 + count2|). The reliability score ranged from zero (unreliable) to one (reliable). Crowd-workers who reported no flowers on one sheet and non-zero number of flowers on the duplicate sheet were assigned a reliability score of zero and were excluded from the analysis (following Park et al. 2018; Peng et al. 2024). We also excluded crowd-workers who recorded the same flower count for all specimens within a set, as this lack of variability suggested inattention to the task. Each herbarium specimen was scored independently by at least three crowd-workers. The median flower count across these crowd-workers was calculated for each specimen and then rounded to the nearest integer. We excluded specimens and observational records in which the number of flowers was zero, leaving 1,120 specimens with records of the number of open flowers on a particular date in the final dataset (median sample size = 24; SD = 13).

### Models used for predicting monthly snow density from climatic variables

We used three regression algorithms to model monthly snow density as a function of temperature, precipitation, and temperature range: generalized additive models (GAMs), random forests (RFs) and generalized linear models (GLMs). The climatic dataset from Prather et al. (2023) was randomly divided into 80% training and 20% testing subsets, and this procedure was repeated 99 times. For each algorithm, we calculated the *R^2^* for each replicate and used the median *R^2^* for comparisons. All three algorithms (RF, GAM and GLM) showed similar model performance in predicting snow density from mean monthly temperature, total monthly precipitation, and monthly temperature range, with median *R^2^* values of 0.62 for RF, 0.57 for GAM and 0.60 for GLM, respectively. Based on comparison of predicted and observed snow density values, RF showed a greater predictive ability than the other two algorithms, with predictions more closely aligned along the 1:1 line (Figure S3). RF was thus selected for estimation of monthly snow density for herbarium specimens.

To evaluate the reliability of PRISM data in representing the climatic conditions of RMBL observations, we additionally matched PRISM climatic data with RMBL observations and applied RF to predict monthly snow density for each observation. For prediction, the climatic data from PRISM associated with herbarium specimens or RMBL observations were likewise partitioned into 80% training and 20% testing subsets 99 times and models with poor predictive performance (*R^2^* < 0.6) were excluded. This approach allowed us to estimate snow conditions specific to the collection date and location of each specimen, providing a more consistent climatic context for comparison with observational records.

### Sensitivity Analysis

Since flowering DOY measurements across consecutive years for the same individuals are not independent, we adopted two alternative approaches to examine whether non-independence among repeated observations affected the results. First, for RMBL observations, we calculated the mean flowering DOY for each species in each year, while each herbarium specimen was treated as an independent record. The model structures were the same as those described above. Second, based on the original model structure, we additionally included random slopes for the year × dataset type interaction to account for species-specific temporal trends and control for the potential influence of interannual variation on flowering phenology.

Because the LMMs assessed only the mean flowering DOY, we complemented these analyses with species-specific quantile regressions (Koenker et al. 2021) to investigate how environmental factors influence early (10^th^ quantile), median (50^th^ quantile), and late (90^th^ quantile) flowering times in both RMBL and herbarium observations. Specifically, each model regressed all observed DOY of a given species against spring mean temperature (or snow density in May) and the number of flowers across years. We extracted species-specific slopes from the 10th, 50th, and 90th quantile regressions based on herbarium specimens and compared them separately with the slopes from the 50th quantile regressions based on RMBL observations.

---

**Table S1.** Monthly climatic variables for RMBL plots obtained from Prather et al. (2023).

| Variable | Description |
| --- | --- |
| Average temperature | Average temperature each month |
| Minimum temperature | Minimum temperature each month |
| Maximum temperature | Maximum temperature each month |
| Mean temperature delta | Difference in mean temperature over the two preceding years for given month |
| Snow density | Cumulative snow density, or snow water equivalent during months of snow |
| Precipitation | Cumulative precipitation each month |
| Growing degree days | Cumulative sum of the daily average temperature each day across each month post snowmelt when the average temperature is $\geq 0^{\circ}\text{C}$ |
| Average discharge rate | Average monthly discharge rate of the East River |
| Average day length | Average number of hours of sunlight each month |
| Mean wind speed | Average modeled wind speed across each month |
| Mean soil moisture % 8in deep | Monthly mean soil moisture % 8 inches below surface level for each month |
| Mean soil moisture % 20in deep | Monthly mean soil moisture % 20 inches below surface level for each month |
| Mean soil temperature % 8in deep | Monthly mean soil temperature % 8 inches below surface level for each month |
| Mean soil temperature % 20in deep | Monthly mean soil temperature % 20 inches below surface level for each month |

**Table S2.**
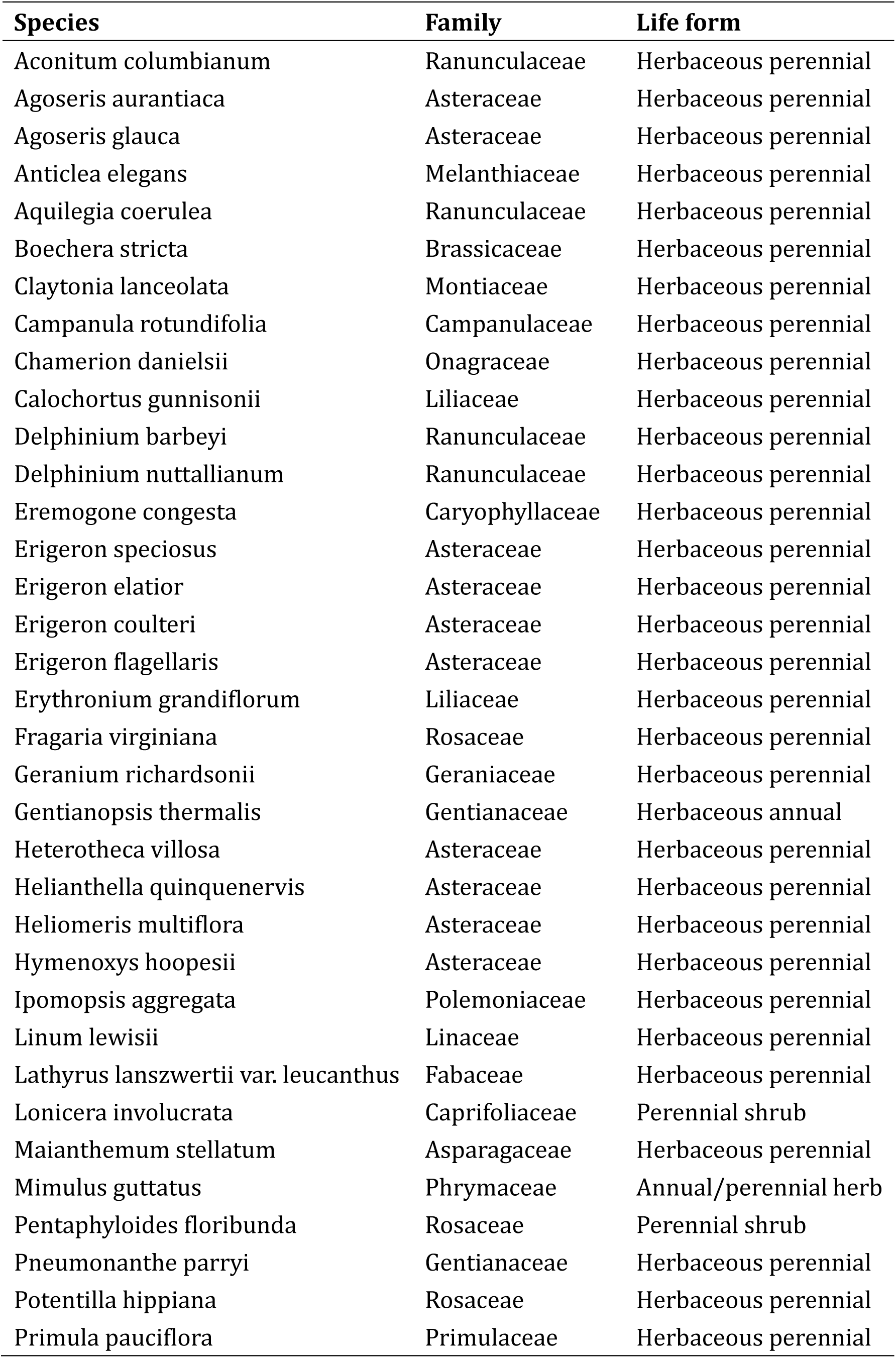
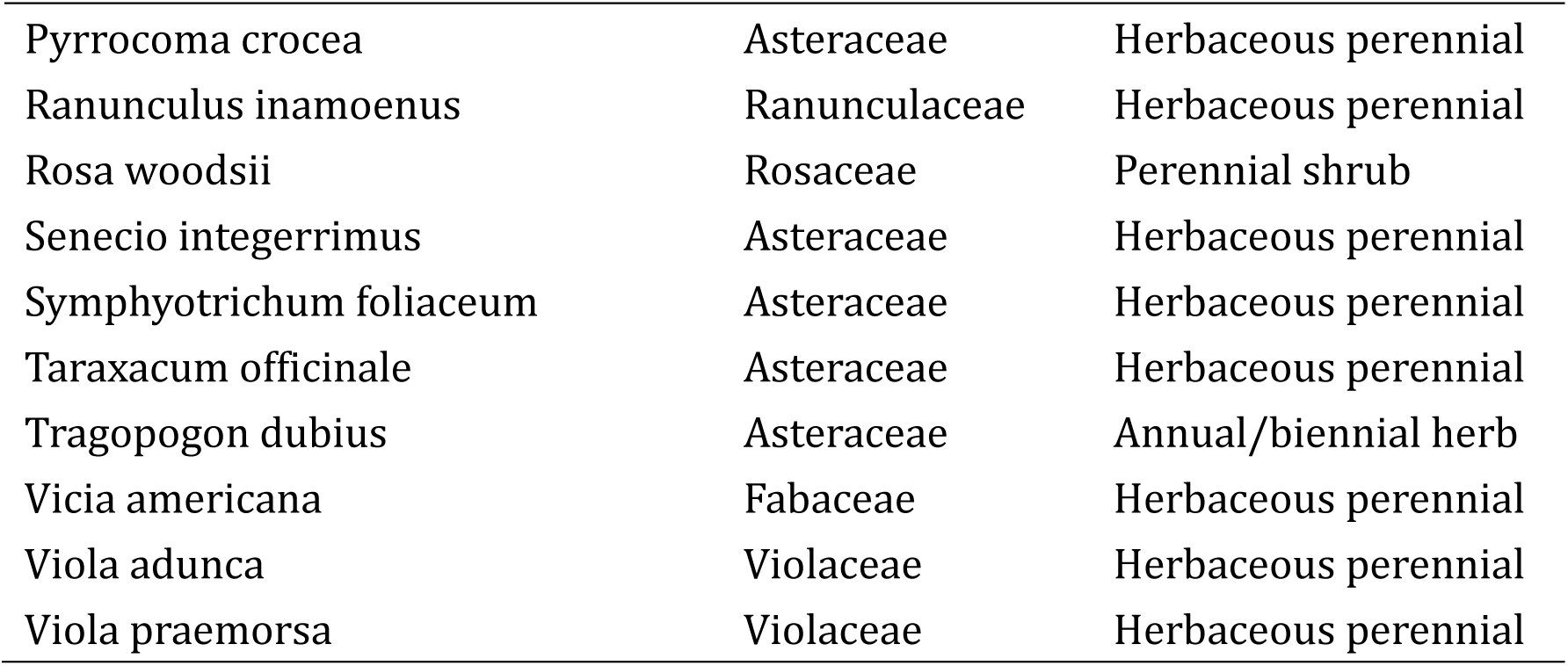
45 plant species used in our study. *Silene antirrhina* was excluded because no specimens fall within the same climatic envelope as the RMBL observation field.

| **Species** | **Family** | **Life form** |
| --- | --- | --- |
| Aconitum columbianum | Ranunculaceae | Herbaceous perennial |
| Agoseris aurantiaca | Asteraceae | Herbaceous perennial |
| Agoseris glauca | Asteraceae | Herbaceous perennial |
| Anticlea elegans | Melanthiaceae | Herbaceous perennial |
| Aquilegia coerulea | Ranunculaceae | Herbaceous perennial |
| Boechera stricta | Brassicaceae | Herbaceous perennial |
| Claytonia lanceolata | Montiaceae | Herbaceous perennial |
| Campanula rotundifolia | Campanulaceae | Herbaceous perennial |
| Chamerion danielsii | Onagraceae | Herbaceous perennial |
| Calochortus gunnisonii | Liliaceae | Herbaceous perennial |
| Delphinium barbeyi | Ranunculaceae | Herbaceous perennial |
| Delphinium nuttallianum | Ranunculaceae | Herbaceous perennial |
| Eremogone congesta | Caryophyllaceae | Herbaceous perennial |
| Erigeron speciosus | Asteraceae | Herbaceous perennial |
| Erigeron elatior | Asteraceae | Herbaceous perennial |
| Erigeron coulteri | Asteraceae | Herbaceous perennial |
| Erigeron flagellaris | Asteraceae | Herbaceous perennial |
| Erythronium grandiflorum | Liliaceae | Herbaceous perennial |
| Fragaria virginiana | Rosaceae | Herbaceous perennial |
| Geranium richardsonii | Geraniaceae | Herbaceous perennial |
| Gentianopsis thermalis | Gentianaceae | Herbaceous annual |
| Heterotheca villosa | Asteraceae | Herbaceous perennial |
| Helianthella quinquenervis | Asteraceae | Herbaceous perennial |
| Heliomeris multiflora | Asteraceae | Herbaceous perennial |
| Hymenoxys hoopesii | Asteraceae | Herbaceous perennial |
| Ipomopsis aggregata | Polemoniaceae | Herbaceous perennial |
| Linum lewisii | Linaceae | Herbaceous perennial |
| Lathyrus lanszwertii var. leucanthus | Fabaceae | Herbaceous perennial |
| Lonicera involucrata | Caprifoliaceae | Perennial shrub |
| Maianthemum stellatum | Asparagaceae | Herbaceous perennial |
| Mimulus guttatus | Phrymaceae | Annual/perennial herb |
| Pentaphyloides floribunda | Rosaceae | Perennial shrub |
| Pneumonanthe parryi | Gentianaceae | Herbaceous perennial |
| Potentilla hippiana | Rosaceae | Herbaceous perennial |
| Primula pauciflora | Primulaceae | Herbaceous perennial |
| *Pyrrocoma crocea* | Asteraceae | Herbaceous perennial |
| *Ranunculus inamoenus* | Ranunculaceae | Herbaceous perennial |
| *Rosa woodsii* | Rosaceae | Perennial shrub |
| *Senecio integerrimus* | Asteraceae | Herbaceous perennial |
| *Symphyotrichum foliaceum* | Asteraceae | Herbaceous perennial |
| *Taraxacum officinale* | Asteraceae | Herbaceous perennial |
| *Tragopogon dubius* | Asteraceae | Annual/biennial herb |
| *Vicia americana* | Fabaceae | Herbaceous perennial |
| *Viola adunca* | Violaceae | Herbaceous perennial |
| *Viola praemorsa* | Violaceae | Herbaceous perennial |

**Table S3.** Importance of seven candidate monthly predictor variables in a random forest model of monthly snow density. Importance values were quantified using percent increase in mean squared error (%IncMSE).

| Variable | %IncMSE |
| --- | --- |
| Mean temperature | 3.792 |
| Total precipitation | 7.097 |
| Mean temperature delta | -1.874 |
| Mean wind speed | -0.622 |
| Temperature range | 6.222 |
| Mean soil temperature 8 inches below surface level | -2.184 |
| Mean soil moisture 8 inches below surface level | 2.976 |

**Table S4.** Shape (*k*) and scale (λ) parameters of the Weibull distribution for each species, estimated separately from RMBL observations and herbarium specimens.

| Species | RMBL observations |  | Herbarium specimens |  |
| --- | --- | --- | --- | --- |
| | Scale ($\lambda$) | Shape ($\kappa$) | Scale ($\lambda$) | Shape ($\kappa$) |
| Aconitum_columbianum | 216.963 | 17.970 | 212.558 | 12.726 |
| Agoseris_aurantiaca | 202.722 | 15.088 | 212.666 | 14.939 |
| Agoseris_glauca | 227.476 | 15.528 | 212.245 | 8.072 |
| Anticlea_elegans | 206.676 | 17.883 | 209.694 | 17.196 |
| Aquilegia_coerulea | 198.148 | 17.194 | 202.418 | 10.943 |
| Boechera_stricta | 194.121 | 7.384 | 207.378 | 14.580 |
| Calochortus_gunnisonii | 213.683 | 18.881 | 207.971 | 30.396 |
| Campanula_rotundifolia | 227.588 | 15.864 | 228.438 | 9.156 |
| Chamerion_danielsii | 227.068 | 26.234 | 218.180 | 21.706 |
| Claytonia_lanceolata | 153.496 | 11.499 | 171.007 | 14.769 |
| Delphinium_barbeyi | 214.861 | 15.932 | 210.499 | 17.666 |
| Delphinium_nuttallianum | 175.752 | 13.517 | 182.312 | 13.943 |
| Eremogone_congesta | 215.423 | 10.578 | 206.586 | 17.778 |
| Erigeron_coulteri | 222.805 | 14.349 | 219.681 | 12.012 |
| Erigeron_elatior | 225.969 | 15.355 | 212.847 | 25.150 |
| Erigeron_flagellaris | 221.913 | 7.405 | 203.853 | 7.709 |
| Erigeron_speciosus | 225.104 | 14.827 | 222.866 | 10.481 |
| Erythronium_grandiflorum | 157.923 | 15.066 | 173.889 | 9.103 |
| Fragaria_virginiana | 179.365 | 10.959 | 202.451 | 7.503 |
| Gentianopsis_thermalis | 238.968 | 16.384 | 232.806 | 13.498 |
| Geranium_richardsonii | 210.859 | 14.301 | 208.132 | 10.617 |
| Helianthella_quinquenervis | 214.285 | 16.792 | 211.611 | 17.032 |
| Heliomeris_multiflora | 237.447 | 18.321 | 221.331 | 12.484 |
| Heterotheca_villosa | 233.338 | 11.905 | 210.526 | 9.147 |
| Hymenoxys_hoopesii | 217.505 | 13.672 | 210.159 | 12.625 |
| Ipomopsis_aggregata | 219.822 | 11.846 | 217.017 | 11.597 |
| Lathyrus_lanszwertii_var._leucanthus | 191.537 | 13.114 | 182.632 | 21.142 |
| Linum_lewisii | 210.27 | 12.451 | 194.075 | 12.482 |
| Lonicera_involucrata | 181.936 | 23.915 | 199.196 | 12.009 |
| Maianthemum_stellatum | 181.266 | 20.849 | 191.494 | 10.533 |
| Mimulus_guttatus | 217.40 | 16.786 | 218.154 | 10.057 |
| Pentaphyloides_floribunda | 212.356 | 13.252 | 209.453 | 9.208 |
| Pneumonanthe_parryi | 242.731 | 21.439 | 236.116 | 15.849 |
| Potentilla_hippiana | 211.763 | 11.522 | 204.212 | 18.684 |
| Primula_pauciflora | 183.513 | 18.144 | 199.932 | 24.912 |
| Pyrrocoma_crocea | 219.586 | 17.355 | 218.040 | 13.459 |
| Ranunculus_inamoenus | 175.086 | 8.907 | 182.658 | 12.286 |
| Rosa_woodsii | 198.474 | 20.142 | 204.737 | 31.191 |
| Senecio_integerrimus | 183.273 | 11.166 | 193.653 | 10.279 |
| Symphyotrichum_foliaceum | 241.019 | 20.625 | 237.555 | 12.623 |
| Taraxacum_officinale | 177.007 | 8.406 | 210.296 | 13.638 |
| Tragopogon_dubius | 228.847 | 14.534 | 208.531 | 9.388 |
| Vicia_americana | 205.103 | 13.712 | 206.172 | 10.425 |
| Viola_adunca | 167.150 | 30.459 | 190.211 | 10.815 |
| Viola_praemorsa | 165.997 | 14.761 | 164.981 | 16.135 |

**Table S5.** Summary of a linear mixed-effect model for comparing the sensitivity of plant flowering day of year (DOY) to snow density in May across three data sources: RMBL field observations with climatic data from Prather et al. (2023), RMBL field observations with PRISM climatic data, and herbarium specimens with PRISM climatic data. Flowering DOY was modeled as a function of snow density, data source, the number of flowers, and the interaction between snow density and data source, all included as fixed effects. For the RMBL observations, we used the maximum number of flowers recorded for each species in each year, whereas for the herbarium specimens, we directly used the number of flowers counted on each individual. Snow density in May and the number of flowers are scaled.

|  | **Estimate** | **SE** | **t value** | **P value** |
| --- | --- | --- | --- | --- |
| Intercept | 201.4 | 3.455 | 58.293 | < 0.001*** |
| Snow density | 3.827 | 0.230 | 16.777 | < 0.001*** |
| Herbarium_PRISM | -3.278 | 4.317 | -0.759 | 0.452 |
| RMBL_PRISM | -2.382 | 0.076 | -31.436 | < 0.001*** |
| The number of flowers | 0.555 | 0.053 | 10.404 | < 0.001*** |
| Snow density: Herbarium_PRISM | -0.573 | 1.081 | -0.530 | 0.599 |
| Snow density: RMBL_PRISM | 0.409 | 0.173 | 2.370 | 0.026* |

\* $P < 0.05$ ; \*\* $P < 0.01$ ; \*\*\* $P < 0.001$

**Table S6.** Summary of a linear mixed-effect model for comparing the sensitivity of plant flowering day of year (DOY) to spring temperature (i.e., mean temperature from April to June) across three data sources: RMBL field observations with local climatic data from Prather et al. (2023), RMBL field observations with PRISM climatic data, and herbarium specimens with PRISM climatic data. Flowering DOY was modeled as a function of spring mean temperature, data source, the number of flowers, and the interaction between spring temperature and data source, all included as fixed effects. For the RMBL observations, we used the maximum number of flowers recorded for each species in each year, whereas for the herbarium specimens, we directly used the number of flowers counted on each individual. Mean spring temperature and the number of flowers are scaled.

|  | Estimate | SE | t value | P value |
| --- | --- | --- | --- | --- |
| Intercept | 201.4 | 3.404 | 59.160 | < 0.001*** |
| Spring temperature | -6.931 | 0.245 | -28.272 | < 0.001*** |
| Herbarium_PRISM | -3.716 | 4.117 | -0.903 | 0.373 |
| RMBL_PRISM | -1.881 | 0.067 | -27.889 | < 0.001*** |
| The number of flowers | 0.697 | 0.047 | 14.738 | < 0.001*** |
| Spring temperature: Herbarium_PRISM | 2.421 | 0.673 | 3.596 | 0.001*** |
| Spring temperature: RMBL_PRISM | 0.557 | 0.169 | 3.302 | 0.003** |

\* $P < 0.05$ ; \*\* $P < 0.01$ ; \*\*\* $P < 0.001$

**Table S7.** Summary of a linear mixed-effect model for comparing the sensitivity of plant mean flowering day of year (DOY) to snow density in May across three data sources: RMBL field observations with climatic data from Prather et al. (2023), RMBL field observations with PRISM climatic data, and herbarium specimens with PRISM climatic data. Mean flowering DOY of each species in each year was calculated for RMBL observations, while each herbarium specimen was treated as an independent record. Mean flowering DOY was modeled as a function of snow density, data source, the number of flowers, and the interaction between snow density and data source, all included as fixed effects. For the RMBL observations, we used the maximum number of flowers recorded for each species in each year, whereas for the herbarium specimens, we directly used the number of flowers counted on each individual.

|  | Estimate | SE | t value | P value |
| --- | --- | --- | --- | --- |
| Intercept | 203.6 | 3.347 | 60.841 | <0.001*** |
| Snow density | 4.342 | 0.205 | 21.23 | < 0.001*** |
| Herbarium_PRISM | -6.136 | 3.997 | -1.535 | 0.134 |
| RMBL_PRISM | -3.001 | 0.172 | -17.402 | < 0.001*** |
| The number of flowers | 0.003 | 0.0007 | 4.62 | <0.001*** |
| Snow density: Herbarium_PRISM | -0.998 | 1.132 | -0.882 | 0.383 |
| Snow density: RMBL_PRISM | -0.519 | 0.173 | -3.008 | 0.003** |

\* $P < 0.05$ ; \*\* $P < 0.01$ ; \*\*\* $P < 0.001$

**Table S8.** Summary of a linear mixed-effect model for comparing the sensitivity of plant mean flowering day of year (DOY) to spring temperature across three data sources: RMBL field observations with climatic data from Prather et al. (2023), RMBL field observations with PRISM climatic data, and herbarium specimens with PRISM climatic data. Mean flowering DOY of each species in each year was calculated for RMBL observations, while each herbarium specimen was treated as an independent record. Mean flowering DOY was modeled as a function of spring temperature, data source, the number of flowers, and the interaction between spring temperature and data source, all included as fixed effects. For the RMBL observations, we used the maximum number of flowers recorded for each species in each year, whereas for the herbarium specimens, we directly used the number of flowers counted on each individual.

|  | Estimate | SE | t value | P value |
| --- | --- | --- | --- | --- |
| Intercept | 203.6 | 3.311 | 61.497 | <0.001*** |
| Mean temperature | -7.424 | 0.242 | -30.669 | < 0.001*** |
| Herbarium_PRISM | -6.343 | 4.066 | -1.560 | 0.128 |
| RMBL_PRISM | -2.071 | 0.139 | -14.868 | < 0.001*** |
| The number of flowers | 0.004 | 0.0006 | 6.077 | < 0.001*** |
| Mean temperature: Herbarium_PRISM | 2.695 | 0.721 | 3.736 | < 0.001*** |
| Mean temperature: RMBL_PRISM | 0.391 | 0.155 | 2.523 | 0.014* |

\* $P < 0.05$ ; \*\* $P < 0.01$ ; \*\*\* $P < 0.001$

**Table S9.** Summary of a linear mixed-effect model for comparing the sensitivity of plant flowering day of year (DOY) to snow density, accounting for the influence of year, across three data sources: RMBL field observations with climatic data from Prather et al. (2023), RMBL field observations with PRISM climatic data, and herbarium specimens with PRISM climatic data. Flowering DOY was modeled as a function of snow density, data source, the number of flowers, and the interaction between snow density and data source, all included as fixed effects. Random slopes for the year × dataset type interaction were also included. For the RMBL observations, we used the maximum number of flowers recorded for each species in each year, whereas for the herbarium specimens, we directly used the number of flowers counted on each individual.

|  | **Estimate** | **SE** | **t value** | **P value** |
| --- | --- | --- | --- | --- |
| Intercept | 201.6 | 3.42 | 58.937 | <0.001*** |
| Mean temperature | 4.413 | 0.233 | 18.969 | < 0.001*** |
| Herbarium_PRISM | -4.636 | 4.329 | -1.071 | 0.291 |
| RMBL_PRISM | -2.423 | 0.075 | -32.386 | < 0.001*** |
| The number of flowers | 0.002 | 0.0002 | 10.829 | < 0.001*** |
| Mean temperature: Herbarium_PRISM | -1.154 | 1.13 | -1.021 | 0.316 |
| Mean temperature: RMBL_PRISM | -0.351 | 0.127 | -2.772 | 0.012* |

**Table S10.** Summary of a linear mixed-effect model for comparing the sensitivity of plant flowering day of year (DOY) to spring temperature, accounting for the influence of year, across three data sources: RMBL field observations with climatic data from Prather et al. (2023), RMBL field observations with PRISM climatic data, and herbarium specimens with PRISM climatic data. Flowering DOY was modeled as a function of spring temperature, data source, the number of flowers, and the interaction between spring temperature and data source, all included as fixed effects. Random slopes for the year × dataset type interaction were also included. For the RMBL observations, we used the maximum number of flowers recorded for each species in each year, whereas for the herbarium specimens, we directly used the number of flowers counted on each individual.

|  | **Estimate** | **SE** | **t value** | **P value** |
| --- | --- | --- | --- | --- |
| Intercept | 201.0 | 3.38 | 59.463 | <0.001*** |
| Mean temperature | -6.866 | 0.252 | -27.248 | < 0.001*** |
| Herbarium_PRISM | -4.104 | 4.152 | -0.988 | 0.329 |
| RMBL_PRISM | -1.887 | 0.068 | -27.561 | < 0.001*** |
| The number of flowers | 0.003 | 0.0004 | 9.389 | < 0.001*** |
| Mean temperature: Herbarium_PRISM | 2.691 | 0.694 | 3.876 | < 0.001*** |
| Mean temperature: RMBL_PRISM | 0.098 | 0.186 | 0.53 | 0.60 |

**Figure S1.**
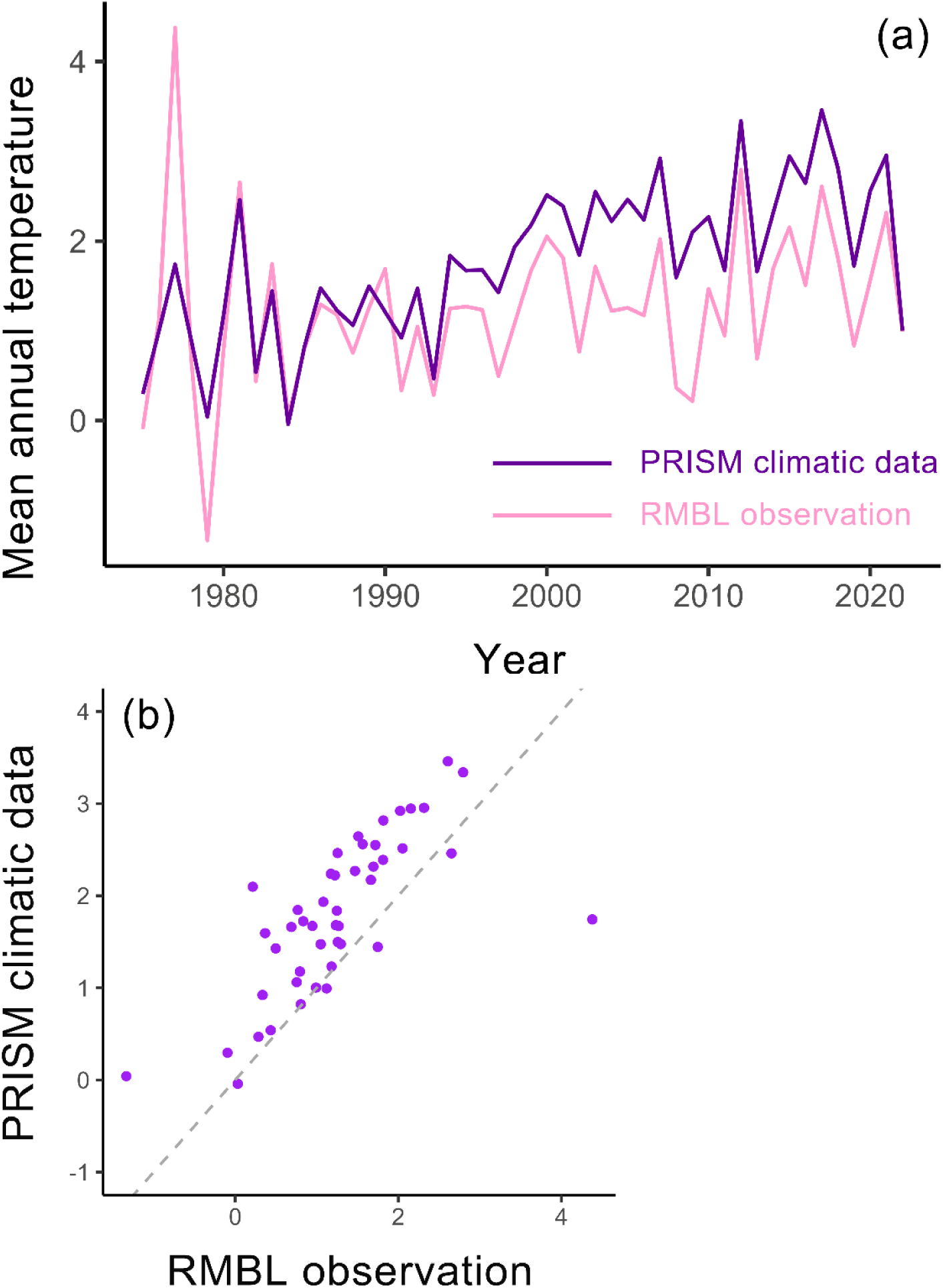
Temporal trend of mean annual temperature in RMBL plots over years, estimated using both PRISM climatic data at a resolution of 800 m (a purple line) and local climatic data from Prather et al. (2023) (a pink line) (a). A pairwise comparison between PRISM climatic data and local climatic data is shown with each point representing a year (b). Grey dashed lines indicate the 1:1 line.

**Figure S2.**
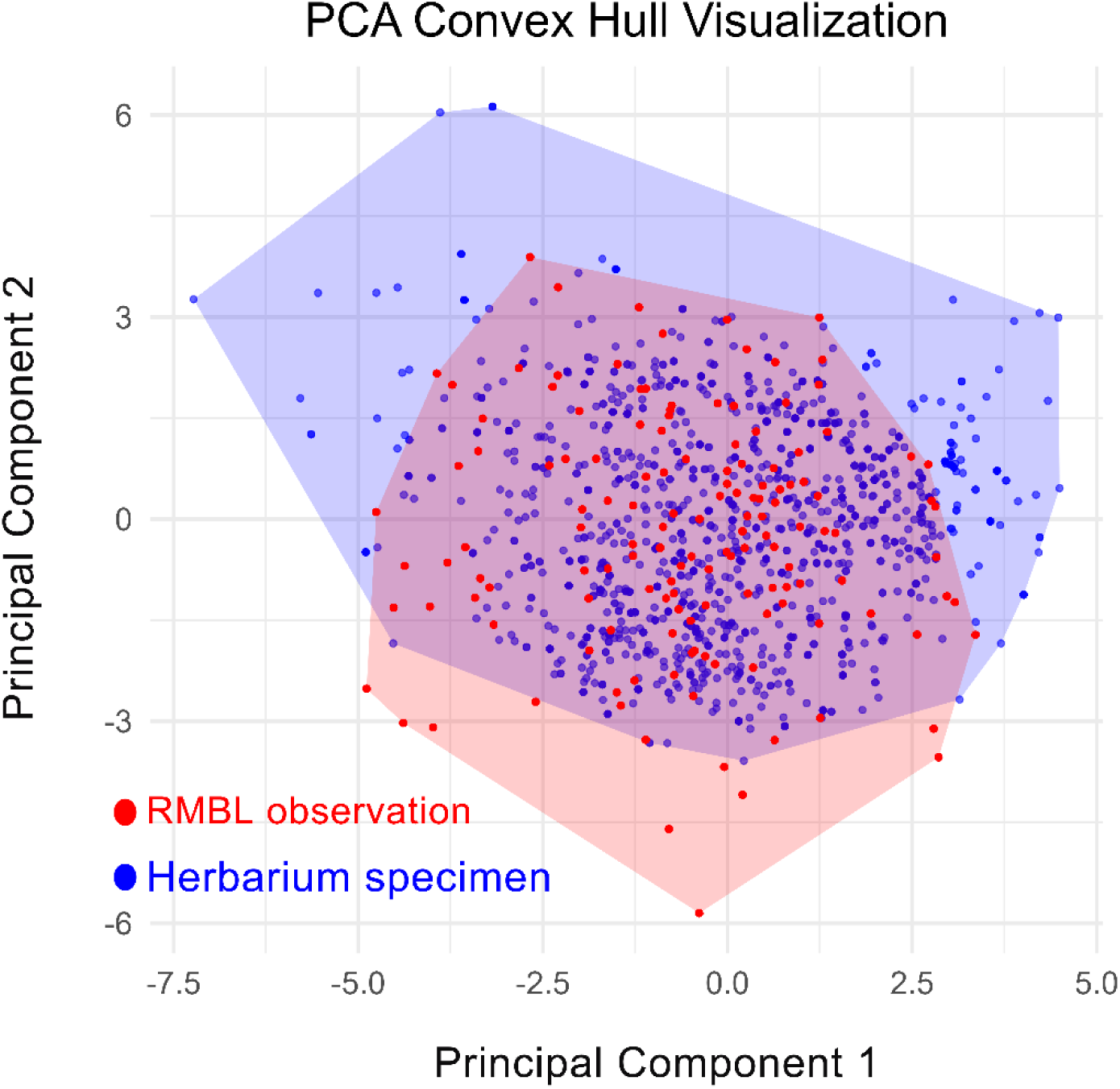
Coverage of the climatic space by herbarium specimens (purple dots and purple convex hull) and RMBL field observations (red dots and convex hull). Climatic space was defined using the first two axes of a principal component analysis (PCA) based on eight bioclimatic variables. Convex hulls represent the full range of climate conditions captured by each dataset across all years. These bioclimatic variables were calculated from monthly temperature and precipitation data from the PRISM dataset.

**Figure S3.**
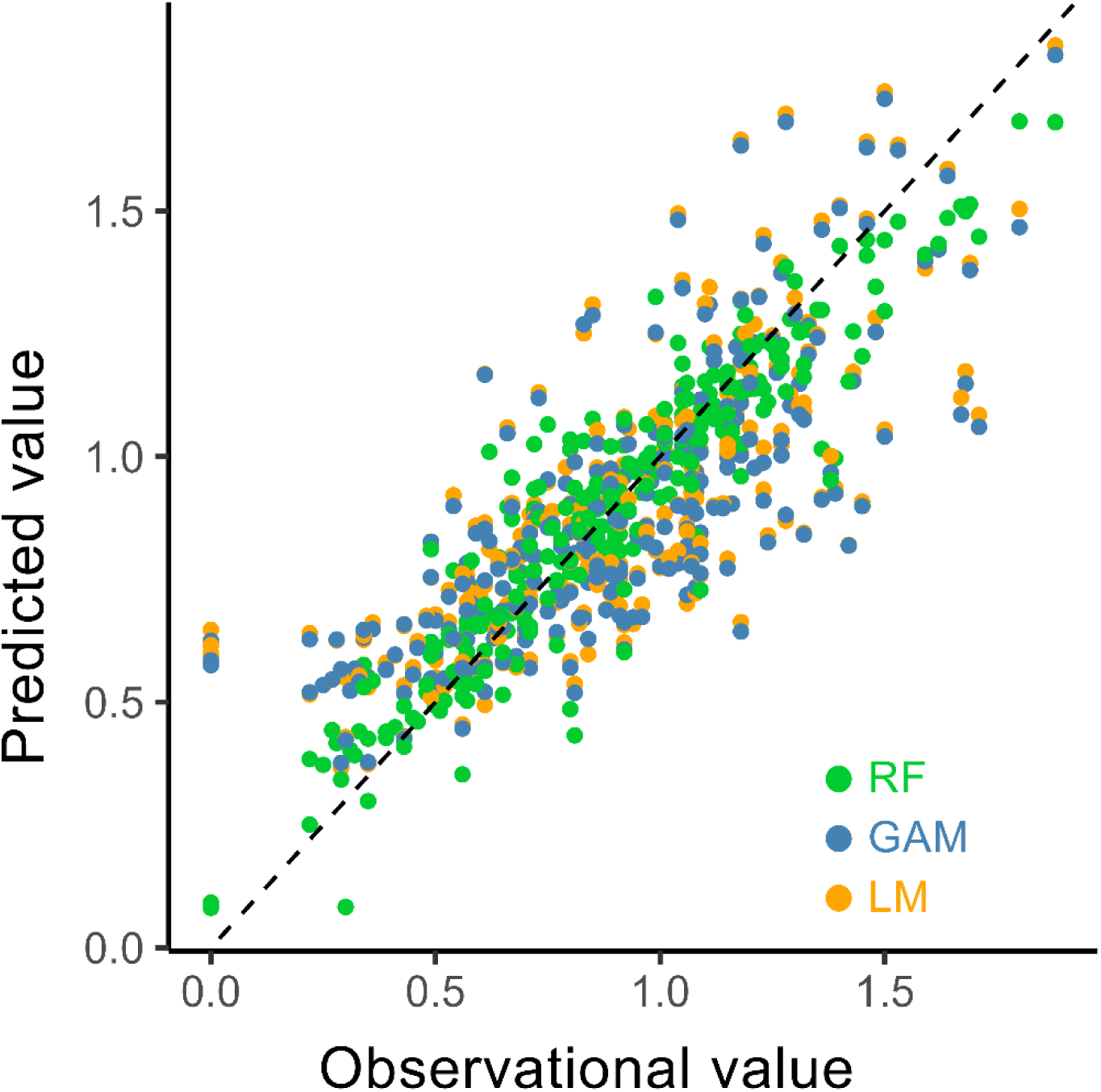
Comparison of predicted versus observed snow density for three fitted algorithms: random forest (RF), generalized additive model (GAM) and linear model (LM).

**Figure S4.**
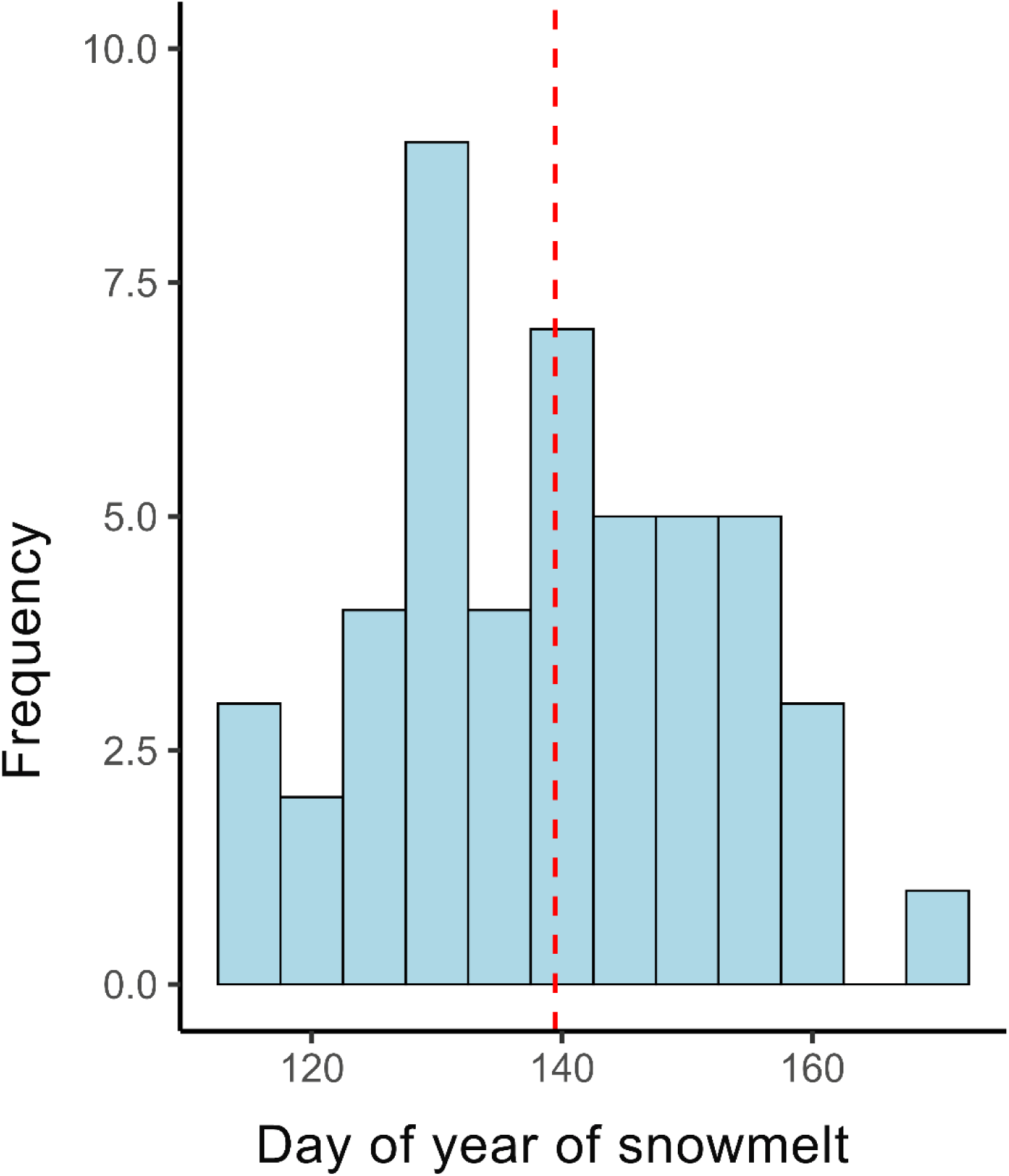
Frequency of snowmelt dates, represented by the day of year (DOY) from 1975 to 2022. The histogram shows the number of years with snowmelt occurring in each DOY class, based on local climatic data from Prather et al. (2023). The vertical line indicates the mean snowmelt date (DOY 139, mid-May).

**Figure S5.**
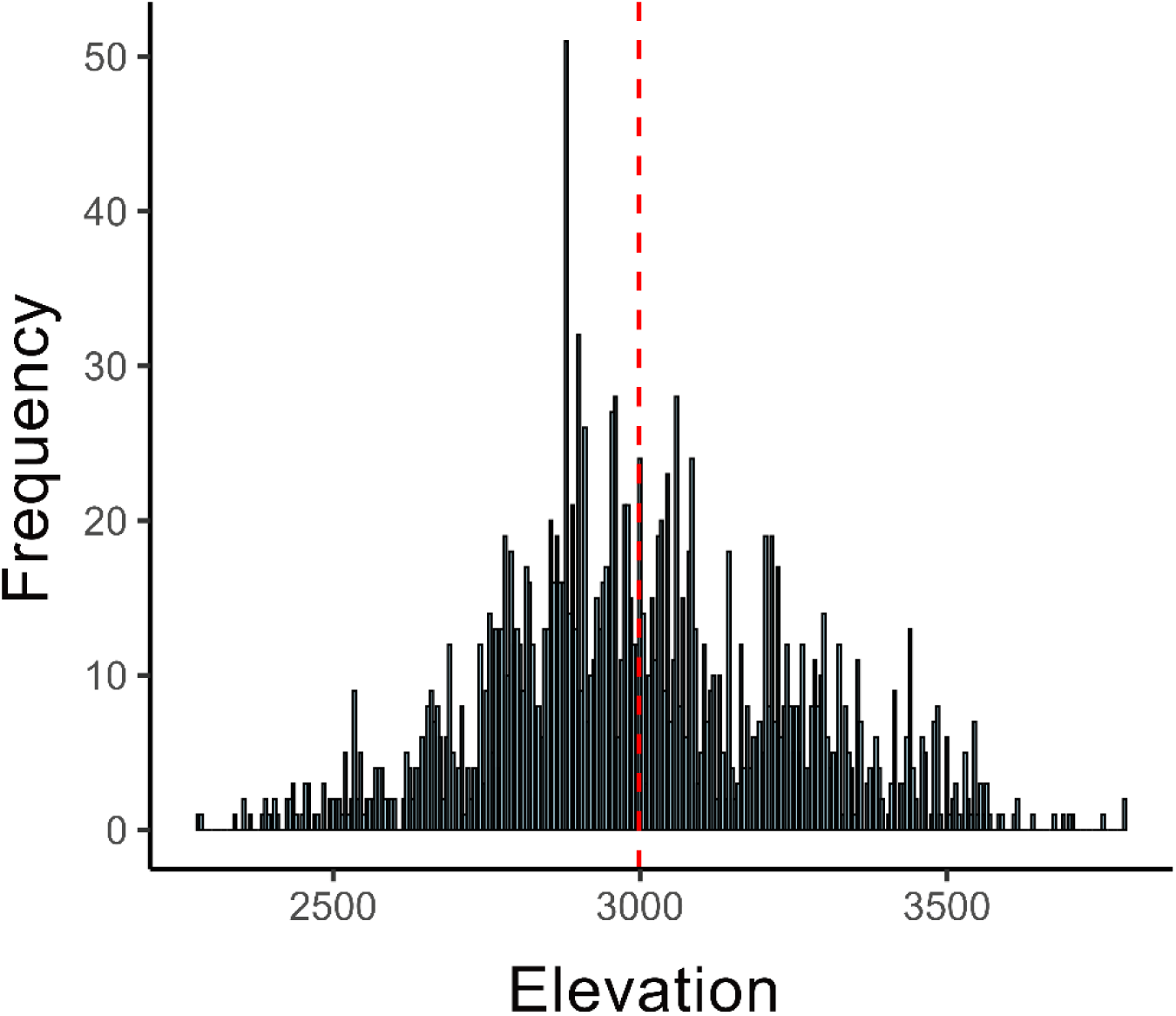
Histogram of elevations of herbarium specimens that fall within the climatic space represented by RMBL plots. Each bar shows the number of specimens within a given elevation class.

